# A conserved homo-dimerization interface in human IFIT1 provides insights into IFIT interactome assembly

**DOI:** 10.1101/152850

**Authors:** Yazan M. Abbas, Saúl Martínez-Montero, Regina Cencic, Jerry Pelletier, Peter D. Pawelek, Masad J. Damha, Bhushan Nagar

## Abstract

The Interferon-Induced Proteins with Tetratricopeptide Repeats (IFITs) are a group of potently expressed Interferon Stimulated Genes that mediate antiviral innate immunity. Previous studies have revealed that most IFITs partake in higher order structures, potentially as part of an ‘IFIT interactome’ that results in viral inhibition. Recent crystal structures of a mutated, monomeric form of IFIT1 revealed the molecular basis of how it recognizes non-self, capped viral mRNAs to selectively inhibit their translation. However, wild-type IFIT1 forms dimers in solution and the role of dimerization was not examined in detail. Here we present a structural and biochemical analysis of wild-type IFIT1 in complex with capped and uncapped RNA. Wild-type IFIT1 forms an antiparallel, elongated dimer that is in stark contrast to the domain-swapped, parallel dimer found in IFIT2. Dimerization takes place through a small, C-terminal interface that is evolutionarily conserved in IFIT1 and IFIT1B proteins. The interface is modular and can be grafted onto IFIT5, which is natively monomeric, to induce dimerization. Mutational analysis of this interface showed that homo-dimerization is not required for full RNA binding or translational inhibition by IFIT1. Sedimentation velocity analytical ultracentrifugation measurements demonstrated a reversible monomer-dimer equilibrium, suggesting that dimerization is of low affinity and could play a role under physiological concentrations, possibly in regulating IFIT interactome assembly. Finally, conformational changes in IFIT1 that occur upon RNA binding provide insight into how RNA enters its binding site in solution.

## Introduction

Interferon (IFN) stimulated genes (ISGs) are a broad class of genes whose expression is triggered downstream of virally activated signaling pathways (1). They number in the hundreds and altogether mediate diverse antiviral effects to promote an antiviral state within infected and uninfected cells. The IFN-induced proteins with tetratricopeptide repeats (IFITs) are among the most potently induced ISGs (2). They are conserved throughout vertebrate evolution, and in humans and most mammals, consist of 5 paralogues: IFIT1, IFIT1B, IFIT2, IFIT3, and IFIT5 (3). IFITs are cytosolic proteins composed of multiple tandem copies of the tetratricopeptide repeat (TPR), a helix-turn-helix motif with the propensity to assemble into super-helical arrays (4). Crystal structures of several IFITs revealed that their TPRs coalesce into three small, super-helical subdomains that come together to form clamp-shaped structures (5-9). TPRs generally mediate protein-protein interactions (10), through which IFITs were previously implicated in modulating several biological processes including cell proliferation (11, 12), migration (13, 14), apoptosis (15), and cytokine signaling (16-18), although the molecular details of these are poorly defined.

However, more recently it was discovered that the primary mode of IFIT action is to deter viral replication through direct interaction with the 5′ ends of viral RNAs (19). This role is best characterized in the ability of IFIT1 and IFIT1B to compete with the eukaryotic initiation factor 4F (eIF4F) for binding to the *N7*-methylguanosine triphosphate (m7Gppp-) cap and cap-proximal nucleotides at the 5′ end of viral mRNAs (6, 20-23). In doing so, IFIT1 and IFIT1B inhibit viral mRNA translation, limiting viral protein production. Endogenous mRNA is protected from IFIT1 and IFIT1B translational inhibition through ‘self’ markers such as ribose 2′-O methylation on the first and sometimes second cap proximal nucleotides (N1 and N2, where N is any nucleotide), which interfere with IFIT1 and IFIT1B binding (6, 20-22, 24, 25). Thus, IFITs play a critical role in antiviral innate immunity by discerning self from non-self. As such, in an attempt to evade IFIT1 and IFIT1B activity, many viruses have acquired the means to produce mRNA displaying 2′-O methylation at N1 (26).

IFIT proteins also target other RNAs as part of their antiviral program. IFIT1 and IFIT5 can recognize uncapped 5′ triphosphate RNA (PPP-RNA) to interfere with the replication of some negative-sense single-stranded RNA viruses (5, 19), and IFIT2 has been shown to interact with AU-rich RNA (8). Conversely, IFIT3 is incapable of binding RNA on its own (5, 19). Thus, by encoding different RNA binding properties and targeting different virus-derived RNAs, IFIT proteins are thought to mediate non-redundant antiviral activities, which is supported by the distinct and context-dependent induction pattern for each IFIT gene (2, 27). However, the expression of multiple IFITs simultaneously in many cell types, and the recent discovery of an IFN-dependent IFIT ‘interactome’ made up of IFIT1, IFIT2, IFIT3, and other host RNA-binding factors suggested cooperation amongst the different IFITs resulting in synergistic or complementary activities (19). At the core of this interactome, IFIT proteins interact directly with each other to form homo- and hetero-oligomers (19), but the nature of these interactions and how they regulate interactome assembly is not clear. Furthermore, how these interactions impact IFIT-RNA binding is not known.

Crystal structures of a mutated form of full-length IFIT1 with capped and uncapped RNA showed that it engages the 5′ end of only single-stranded RNA through a narrow, positively-charged RNA-binding tunnel at the center of the protein (6). The crystal structure of IFIT5 in complex with PPP-RNA showed a similar RNA-binding tunnel (5), which recognizes only uncapped RNA due to protein residues at one end of the tunnel blocking any progression beyond the PPP moiety (5, 6). Notably, biochemical and structural analyses of IFIT5 revealed that it exists exclusively in a monomeric state in solution that undergoes a conformational change upon RNA binding (5). Prior to RNA binding, IFIT5 appears to exist in a more open, discrete conformation in solution that compacts and wraps itself around the incoming RNA. Two extended, non-TPR α-helices, α15 and α16, form a pivot region in IFIT5 that regulate the closure of the protein around the RNA. On the other hand, the crystal structure of RNA-free IFIT2 showed that it formed parallel domain-swapped dimers mediated by the exchange of three α-helices (α7- α9) from its second subdomain, resulting in two adjacent RNA-binding channels with relatively flexible C-terminal regions (8). The crystal structures of an N-terminal fragment of IFIT1 and RNA-bound IFIT1 showed no evidence for domain swapping and therefore how IFIT1 forms higher order structures and the nature of RNA-dependent conformational changes in IFIT1 are as yet unclear (5, 6).

We report here crystal structures of wild-type, human IFIT1 bound to capped and uncapped RNAs. The structures show that IFIT1 forms an antiparallel, end-to-end, S-shaped dimer using its extreme C-terminus to form a pseudo-continuous TPR super-helix, in stark contrast to the globular, domain-swapped, parallel dimer of IFIT2. Biophysical and mutational analysis confirm the presence of IFIT1 dimers in solution, and combined with functional assays, suggest that homo-dimerization is not required for full RNA binding and translational inhibition. Finally, comparison of the different RNA-bound forms of wild-type IFIT1, complemented with limited proteolysis, suggests a role for the C-terminal subdomain and an mRNA ‘cap-binding loop’ in mediating conformational changes that have implications for its antiviral activity. Our work uncovers a novel dimerization mechanism within the IFIT family and suggests a model for mRNA binding in solution.

## Results

### IFIT1 forms dimers in solution

IFIT1 has been reported to oligomerize and bind RNA with positive cooperativity (22), but the stoichiometry and mechanism of oligomerization are unknown. Therefore, we first wanted to validate the IFIT1 molecular weight (MW) in solution by subjecting it to multi-angle light scattering with inline size-exclusion chromatography (SEC-MALS, **Fig. 1A**). IFIT1 was injected into SEC-MALS at 180 μM (10 mg/ml), and the protein migrated predominantly as a single peak with an estimated MW of 101.4 kDa, corresponding to an IFIT1 dimer (expected dimer MW 111 kDa). Interestingly, the elution profiles consistently displayed a non-symmetric Gaussian shape with a tail on the low MW side, suggesting that the dimeric state may be in a dynamic equilibrium.

To better understand the potential dynamic nature of dimerization in solution, we turned to sedimentation velocity analytical ultracentrifugation (AUC, **Fig. 1B** and **Table S1**), performed at a range of temperatures (12-24 °C) and sample concentrations (3.3, 8.5 and 18.0 μM). SEDFIT *c*(*S*) analysis of sedimentation velocity data collected at 16 °C revealed the presence of a major sedimenting species in the IFIT1 preparation that exhibited a concentration-dependent increase in sedimentation coefficient (**Fig. 1B**). Such a concentration-dependent change in sedimentation coefficient is consistent with a reversible monomer-dimer interaction occurring during the run (28). Furthermore, the observed major *c*(*S*) peak was observed to narrow as a function of concentration, being broadest at the lowest concentration measured, consistent with an overall change in shape of the sedimenting species. Similar concentration-dependent increases in *S*20,W were observed between 12 °C and 24 °C (**Table S1**, column 3). The apparent MW of the major sedimenting species calculated by SEDFIT varied between 85-90 kDa, between the monomer MW (55.5 kDa) and dimer MW (111 kDa), as expected for a rapidly reversible monomer-dimer equilibrium occurring during the sedimentation velocity run. That the apparent MW of the major sedimenting species is closer to the predicted dimer MW, suggests that the equilibrium position is towards dimer formation at all concentrations examined. At the lowest concentration, the major sedimenting species accounted for roughly ~75 % of the total optical density at 280 nm, which increased to ~95 % at the highest concentration (**Table S1**, column 4).

**Figure 1.**
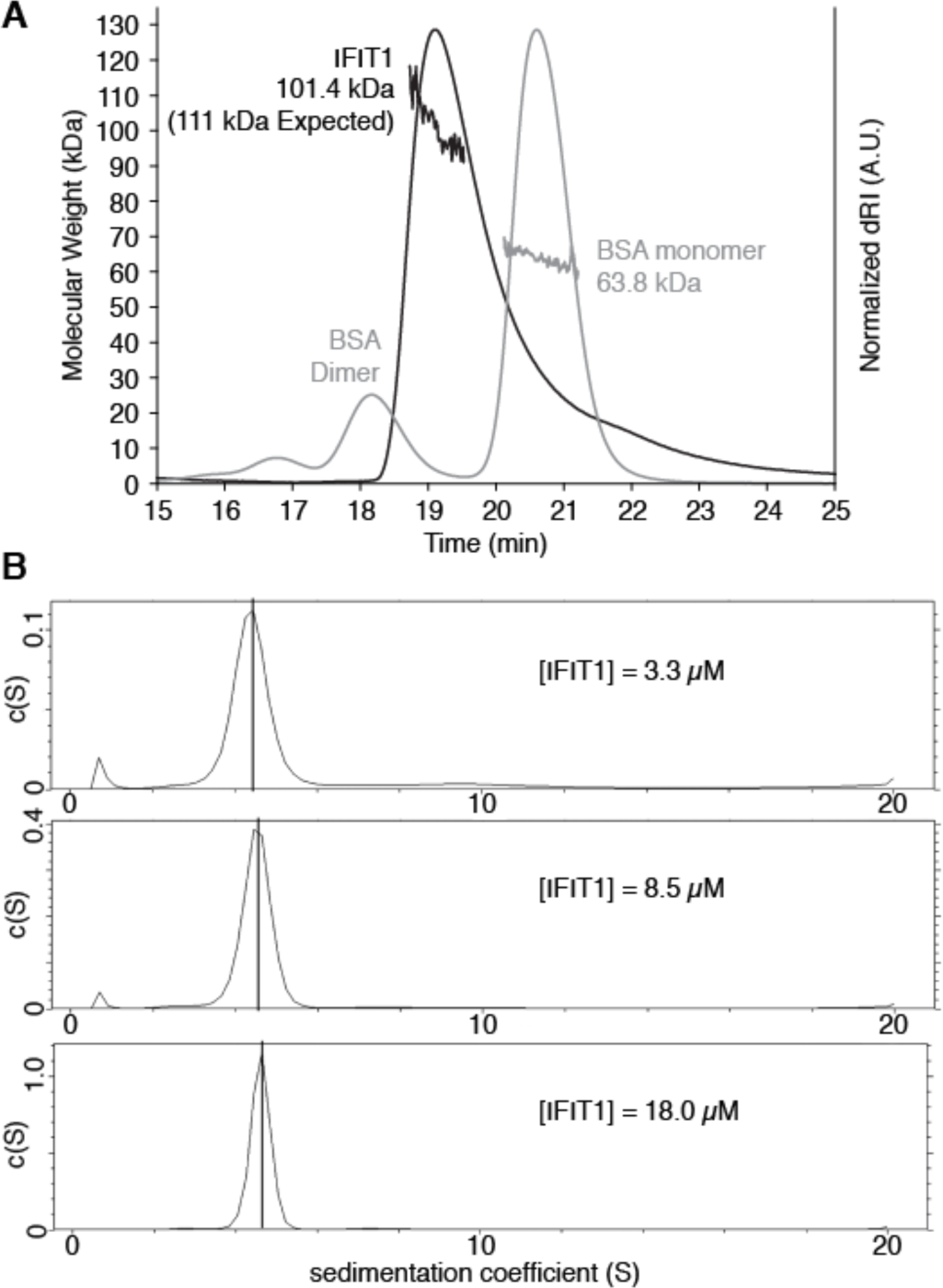
Human IFIT1 forms dimers in solution. **(A)** SEC-MALS of human IFIT1 injected at 10 mg/ml on a Superdex 200 Increase 10/300 column. Shown are the estimated molecular weight and normalized differential refractive index (dRI) as a function of time. BSA (10.5 mg/ml) was performed as a control. **(B)** Sedimentation velocity analytical ultracentrifugation of IFIT1. Sedimentation data were collected at 16 °C and analyzed using SEDFIT. Upper panel: c(S) distribution of IFIT1 at 3.3 μM; middle panel: c(S) distribution of IFIT1 at 8.5 μM; lower panel: c(S) distribution of IFIT1 at 18.0 μM. Vertical lines in peaks correspond to untransformed sedimentation coefficients (4.40 S at 3.3 μM, 4.52 S at 8.5 μM, and 4.61 S at 18.0 μM) calculated by peak integration of c(S) distribution using SEDFIT. Sedimentation coefficients transformed to *S*_20,W_ values can be found in **Table S1**.

### Crystallization and structure determination of human IFIT1

To gain insight into the mechanism of IFIT1 dimerization, we crystallized full-length human IFIT1 (residues 1-478) in complex with PPP- and m7Gppp- containing oligoadenylates (oligoA, four nucleotides in length). RNA-bound IFIT1 crystallized as a dimer with two molecules in the asymmetric unit (ASU), and the crystals diffracted X-rays to resolutions of 2.65 Å (PPP-oligoA), and 2.8 Å (m7Gppp-oligoA). The initial structures were solved by molecular replacement using RNA-bound IFIT5 as a search model, and subsequent structures determined by rigid body refinement (see **Table S2** for data collection and refinement statistics and **Fig. S1 and Fig. S2** for representative protein and RNA electron density, respectively). Superposition of all four protein chains from the two co-crystal structures yields backbone root mean square deviations (r.m.s.d.) of 0.2-0.5 Å for all C_α_ atoms, indicating minimal differences between the two molecules of the ASU or between the different RNA-bound forms (**Fig. S3A**).

We also determined a high resolution, 1.58 Å structure of an RNA-bound, monomeric variant of IFIT1, generated with the aid of the IFIT1 dimer structure determined here (discussed below). The structure of each molecule of IFIT1 from the wild-type dimer form is similar to the monomeric variant, except for small changes at flexible regions and near the protein C-terminus that are likely due to differences in crystal packing (**Fig. S3, B and C**). The details of IFIT1-RNA binding have already been described in an earlier study (6), and here we focus mainly on the mechanism of dimerization and conformational changes associated with RNA binding.

### Overall structure of the RNA-bound human IFIT1 dimer

The two molecules of the ASU interact in a tail-to-tail manner exclusively through their C-terminal regions to form a near two-fold symmetric, S-shaped dimer with ~173° rotation between them and approximate dimensions of 140 Å x 80 Å x 50 Å (**Fig. 2A**). Each monomer of IFIT1 is composed of 23 α-helices, 18 originating from its 9 TPR motifs, organized into 3 distinct subdomains (**Fig. 2A**). As with IFIT5, the subdomains come together to form a positively-charged RNA binding tunnel at the core of the protein, which in IFIT1 encircles the cap and four RNA nucleotides (**Fig. 2, A, B and C**) giving rise to the preference for binding only ssRNA with unstructured 5′ ends or dsRNA with 5′ overhangs (5, 6, 23). The m7G moiety of the cap sits at one end of the tunnel inside a relatively hydrophobic cap-binding pocket between subdomains I and II (**Fig. 2A, B and D**). The m7G electron density in the 2.8 Å model is not well defined (**Fig. S2**), consistent with the presence of two base conformations (*syn* and *anti*) observed in the 1.58 Å structure of monomeric IFIT1 with m7Gppp-RNA (6). As before, one side of the m7G base interacts with Trp 147 (through π-π stacking) and Ile 183, while the other contacts Leu 46 and Thr 48 from a ‘cap-binding loop’ (**Fig. 2D**).

**Figure 2.**
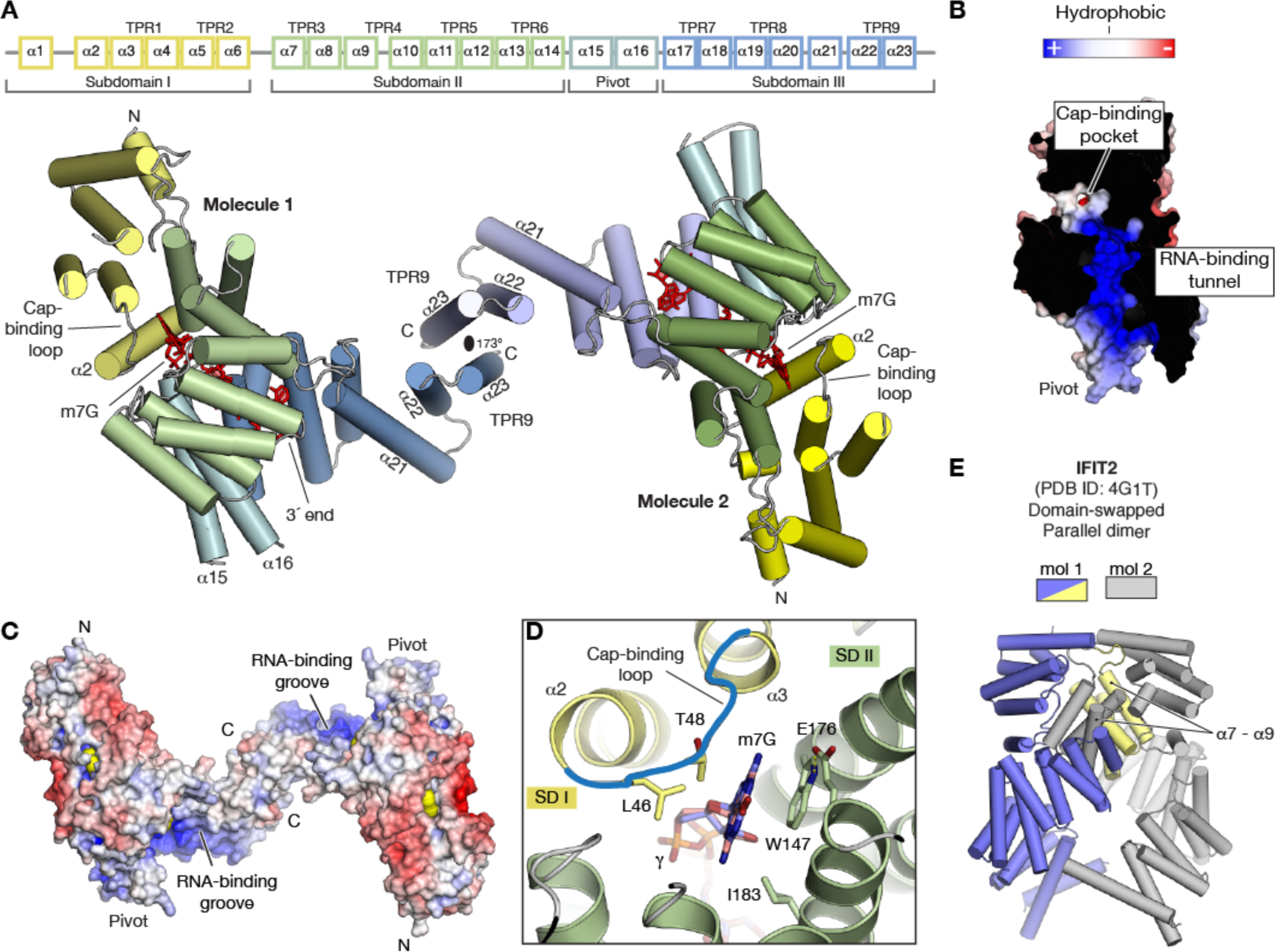
Overall structure of RNA-bound human IFIT1 dimer. **(A)** Schematic of human IFIT1 subdomains and cartoon representation of the overall structure colored by subdomain. **(B)** Cross section of the IFIT1 RNA-binding tunnel and **(C)** IFIT1 dimers colored by surface electrostatic potential from negative (-10 kTe^-1^; red) to positive (+10 kTe^-1^; blue). The 3′ end of the RNA exits from the RNA-binding tunnel and points towards an RNA-binding groove at the C-terminus of each molecule. **(D)** Close-up of the cap-binding pocket with m7G interacting residues represented as sticks and colored according to subdomain. **(E)** In IFIT2, dimerization is through domain-swapping by exchanging helices α7-α9 (pale yellow and grey) from subdomain II between the two molecules.

The RNA-binding tunnel continues along the central super-helix of the protein, spanning subdomain II (TPR3-TPR6), the pivot region (α15+α16), and the N-terminal portion of subdomain III (TPR7-TPR8) (**Fig. 2A**). The 3′ end of the RNA (N4) emerges from the tunnel and points towards a positively-charged, solvent exposed groove which may contribute to non-specific RNA binding (**Fig. 2C**) (6). As with IFIT5, α21 intervenes between TPR8 and TPR9 to cap the central super-helix and alter the right-handed super-helical trajectory at the end of each molecule (**Fig. 2A**). TPR9, in turn, mediates IFIT1 dimerization. The overall arrangement of IFIT1 dimers thus positions the two RNA-binding tunnels in a roughly antiparallel fashion with respect to one another, and results in two non-contiguous RNA-binding surfaces (**Fig. 2, A and C**).

The elongated IFIT1 dimer is distinct from that of IFIT2 (44 % sequence identity), which forms a parallel side-to-side dimer through N-terminal domain swapping (**Fig. 2E**) (8). However, the structure of individual IFIT1 monomers is very similar to IFIT2 and IFIT5 with backbone r.m.s.d’s of 2-4 Å (**Fig. S4A**), which improves to 0.44-1.1 Å when only subdomains I, II, or III are used in the superposition (**Fig. S4B**).

### Mechanism of IFIT1 dimerization

TPR motifs are generally composed of a degenerate, 34 amino-acid sequence that folds into a pair of antiparallel helices (helices A and B) (4). Tandemly arranged TPR motifs will then assemble into super-helical arrays through conserved hydrophobic interactions between helices A and B within one motif, and between one TPR and helix A′ of the following motif. In this manner, the minimal building block for a TPR array is 1.5 TPR motifs (or AB-A′), and the repeating pattern (i.e. AB-A′B′-A″B″) continues until the array is capped at its end by an A-like helix. When present at the C-terminus of a protein, this capping helix also functions as a ‘solubility helix’. For example, in IFIT5, the very C-terminal helix α24 follows TPR9 to bury its hydrophobic surface and prevent further extension of the super-helix (**Fig. 3A**). In comparison, IFIT1 is prematurely truncated because it lacks a corresponding helix α24 (**Fig. 3A and 4A**). The resulting exposed hydrophobic surface on TPR9 is instead capped by an opposing TPR9 from another molecule, thus forming a tail-to-tail elongated dimer.

**Figure 3.**
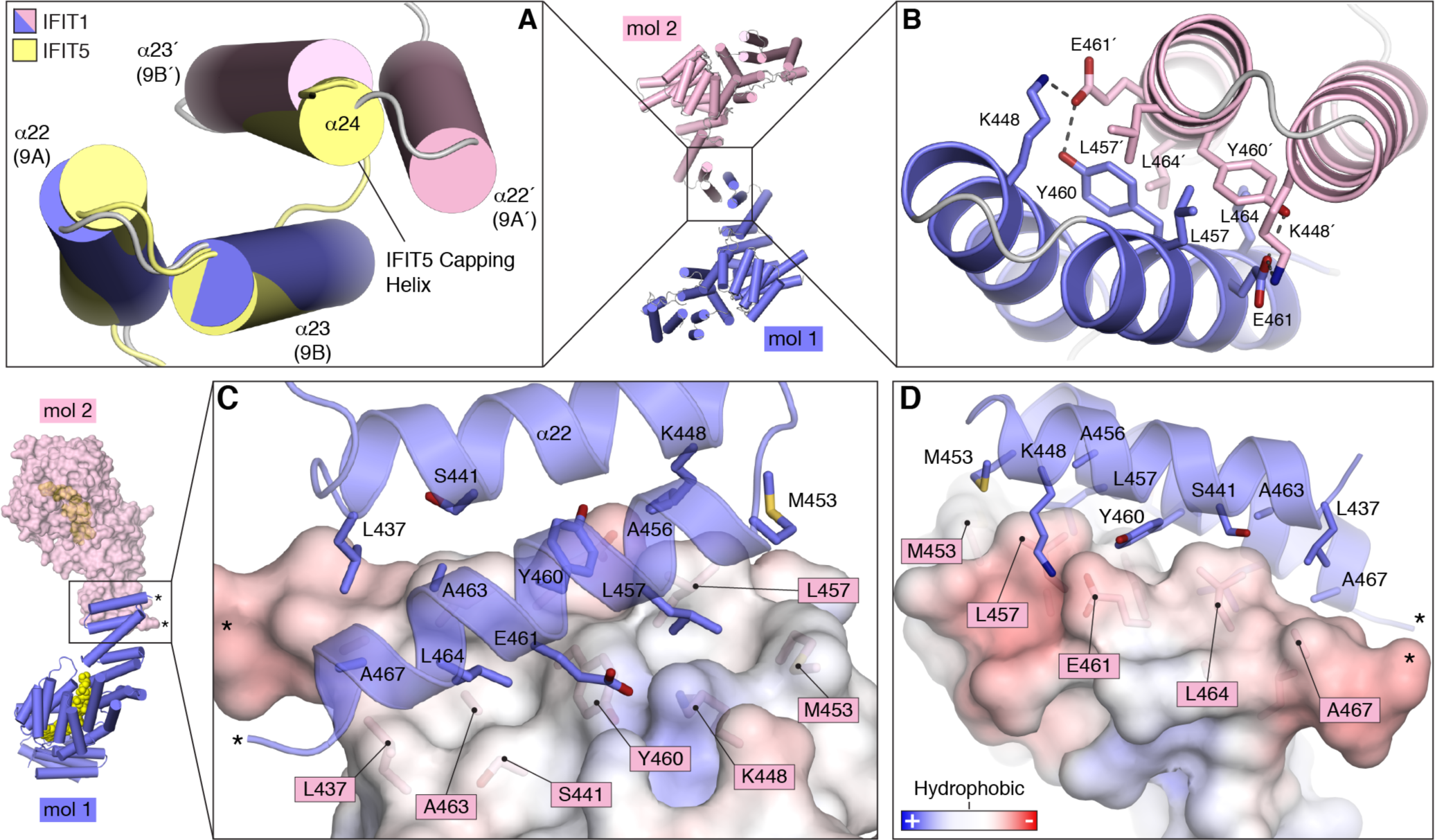
Structural basis for IFIT1 dimerization. **(A)** Close-up of TPR9 (helices 9A and 9B) from human IFIT1 with IFIT5 superposed. Compared to IFIT5, IFIT1 proteins are truncated and lack helix α24, which in IFIT5 functions as a C-terminal capping helix. The two protomers are referred to as IFIT1 (blue) and IFIT1′ (light pink). **(B)** Cartoon/stick representation of the critical residues at the IFIT1 dimerization interface in the same orientation as (A). These residues make up most of the buried surface area between the dimers, and were targeted in the mutational analysis. **(C and D)** Two orthogonal views showing cartoon representation of IFIT1 molecule 1 (blue) and surface representation of IFIT1 molecule 2 colored according to electrostatic potential from -10 kTe^-1^ (red) to + 10 kTe^-1^ (blue). Residues at the interface are shown as sticks. The C-terminus of each molecule is indicated by asterisks for reference.

The dimer interface buries ~ 530 Å^2^ of surface area (determined by PISA, (29)) and is composed of several small and large hydrophobic residues, including Leu 457, Tyr 460, and Leu 464 from each TPR9 motif (**Fig. 3, B, C and D**). The interface is further stabilized by reciprocal hydrogen bonds between Tyr 460 and Glu 461, and salt bridges between Lys 448 and Glu 461 (**Fig. 3B**). The residues involved in dimerization are conserved across mammalian IFIT1 genes, indicating an evolutionarily conserved mechanism for dimerization (**Fig. 4, A and B**). The interface is also conserved in IFIT1B proteins, suggesting that they use the same mechanism of homo-dimerization.

**Figure 4.**
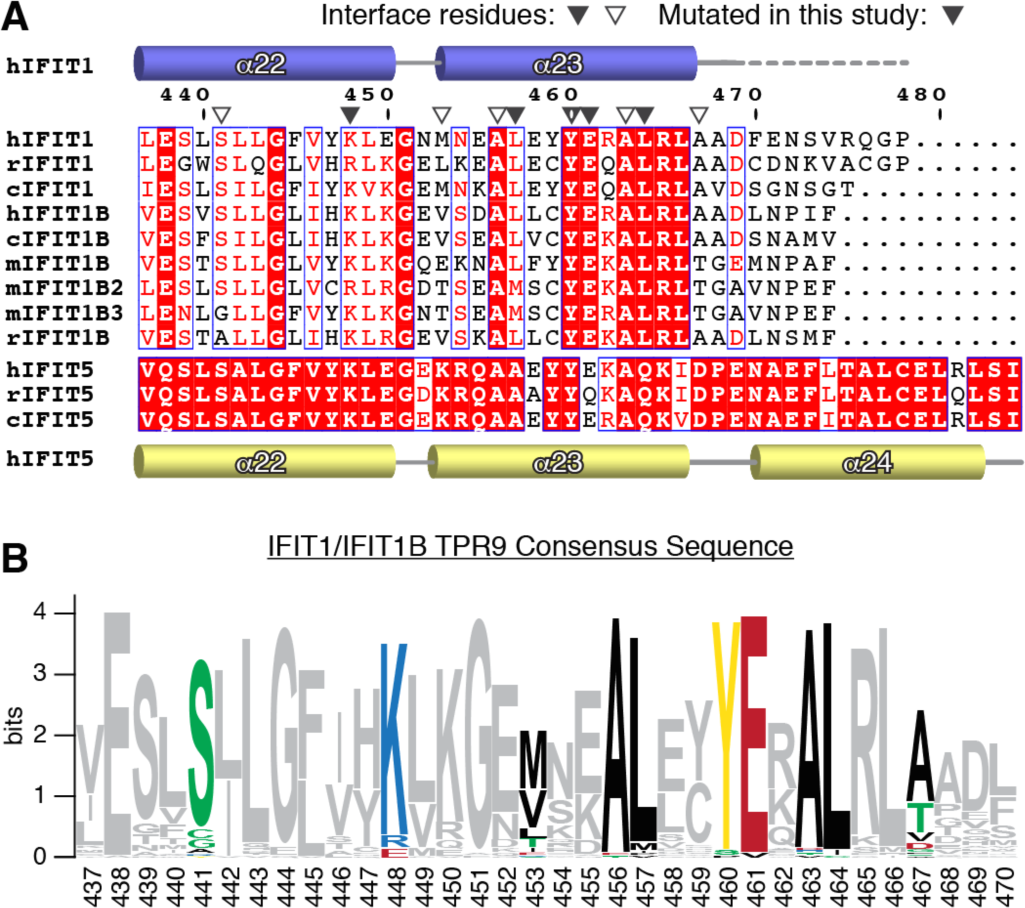
The IFIT1 dimerization interface is conserved in IFIT1 and IFIT1B proteins. **(A)** TPR9 Sequence alignment of select IFIT1, IFIT1B and IFIT5 residues from human, mouse, cat, and rabbit. Secondary structure of IFIT1 and IFIT5 are depicted above and below. The ruler corresponds to human IFIT1 residue numbers. Compared to IFIT1 proteins, IFIT5 proteins are extended and have an additional α-helix (α24). **(B)** WebLogo sequence consensus of 86 IFIT1 and IFITB genes showing only TPR9. Interface residues are colored according to property, and other residues are in gray. The dimerization interface residues are highly conserved.

### Mutational analysis of dimerization

The above described dimer is but one of several distinct dimerization interfaces in the crystal lattice. To confirm that the IFIT1 dimerization observed in our solution experiments is the one mediated by its C-terminus and not one of the other crystal packing interfaces, we performed a mutational analysis of TPR9 which we assayed by analytical gel filtration (**Fig. 5A**). Wild-type and mutant IFIT1 proteins were injected at 2 mg/ml (~ 36 μM) and, as controls, we compared their gel-filtration profiles to that of IFIT2 (109 kDa dimer), IFIT5 (56 kDa monomer), and BSA. The single point mutants L457E, Y460E, and L464E, as well as the double mutants L457A/L464A (IFIT1^DM-A^) and L457E/L464E (IFIT1^DM-E^) all disrupted dimerization. K448E had no impact, indicating that the K448-E461 salt bridge is dispensable, and E461A and E461K mutants could not be expressed. SEC-MALS was repeated to confirm the monomer MW of IFIT1^DM-E^ (**Fig. S5A**), and sedimentation velocity AUC of IFIT1^DM-E^ revealed that at 20 °C the SEDFIT-calculated apparent MW of the major sedimenting species (52.7, 45.1, and 49.1 kDa at 3.3, 8.5 and 18.0 μM, respectively) corresponded to the predicted monomer MW, with no monomer-dimer equilibrium apparent. The mutations do not disrupt the protein fold, since circular dichroism (CD) spectroscopy of IFIT1, IFIT1^DM-A^, and IFIT1^DM-E^ revealed no discernable differences (**Fig. S5B**). Additionally, the crystal structure of RNA-bound IFIT1^DM-E^ described in our previous analysis of IFIT1-RNA interactions revealed no major differences between the wild-type and mutant protein (**Fig. S3B**).

**Figure 5.**
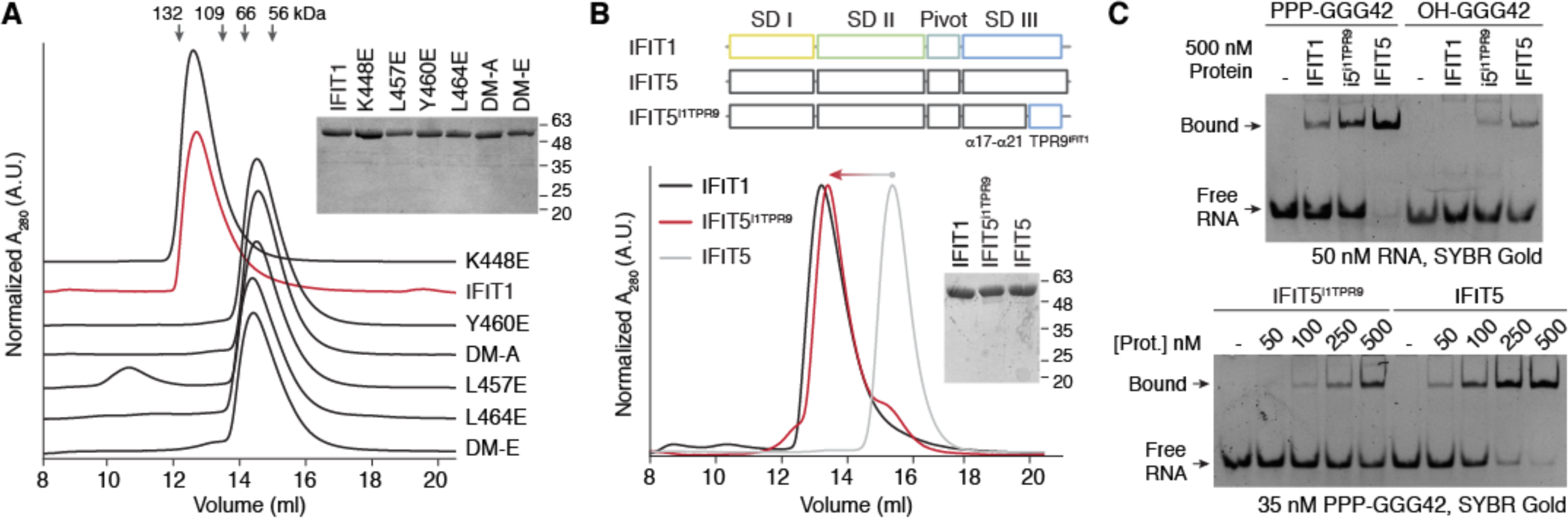
Mutational analysis of IFIT1 dimerization. **(A)** Gel filtration (Superdex 200 Increase 10/300) analysis of IFIT1 mutants (at 2 mg/ml each) with SDS-PAGE of the input proteins (1 μg per lane). Each UV trace was offset along the y-axis for clarity. The peak maximum of IFIT1 wt was arbitrarily set to 1, and each trace shows the peak height relative to IFIT1. The bars above correspond to the migration of BSA dimer (132 kDa), IFIT2 dimer (109 kDa), BSA monomer (66 kDa), and IFIT5 monomer (56 kDa). **(B)** Schematic of the IFIT5-IFIT1 chimera (IFIT5^i1TPR9^), and gel filtration (Superdex 200 Increase 10/300) analysis of the indicated proteins (at 2 mg/ml each) with SDS-PAGE of the input proteins (2 μg per lane). The chimera behaves as a dimer on gel filtration similar to IFIT1. **(C)** SYBR Gold stained electromobility gel-shift assays between a 42-nucleotide PPP-RNA (GGG42) and the indicated proteins. OH-RNA was utilized as a negative control for non-specific RNA binding in the upper panel.

To determine if TPR9 alone is responsible for dimerization, we created a chimera of human IFIT5 α1-α21 (residues 1-435) and human IFIT1 α22-α23 (TPR9, residues 437-478). Analytical gel filtration showed that the chimera can form homo-dimers in solution (**Fig. 5B**), and electro-mobility gel shift assays (EMSAs) between the chimera and a 42-nucleotide RNA (GGG42) shows that it retains binding to PPP-RNA (**Fig. 5C**). However, PPP-RNA binding by the chimera is weaker compared to IFIT5, possibly because dimerization interferes with the dynamics of IFIT5 closure around the RNA. Taken together, the mutational analysis confirms the role of TPR9 and its hydrophobic surface in mediating IFIT1 dimerization, and shows that this motif can behave as an independent dimerization module.

### IFIT1 dimers form a pseudo-continuous TPR super-helix

The central portion of IFIT1 dimers (TPR9s and several preceding helices) where they associate bears a striking resemblance to small TPR domains such as the synthetic TPR protein, cTPR3 (4), made up of three idealized motifs (**Fig. 6A**). In fact, the residues involved in IFIT1 dimerization from the second helix of TPR9 are highly conserved when compared to consensus TPR motif sequences derived from a large database of TPR containing proteins (**Fig. 6C**) (4). These residues are usually critical for mediating the intra-molecular TPR stacking interactions that give rise to continuous TPR arrays (**Fig. 6B**). Interestingly, however, there is one key difference between the inter-molecular stack at the IFIT1 C-terminus and canonical intra-molecular TPR stacking. Whereas canonical TPR stacking is between one TPR and helix A of the next, in IFIT1 the inter-molecular stacking is between TPR9 and helix B of the adjoining TPR9 instead of another A-like helix (**Fig. 6, A and B**). Thus, the two TPR9 motifs of each IFIT1 molecule interact in a manner that is analogous to intra-molecular TPR stacking, and form a pseudo-continuous TPR super-helix which is responsible for IFIT1 dimerization.

**Figure 6.**
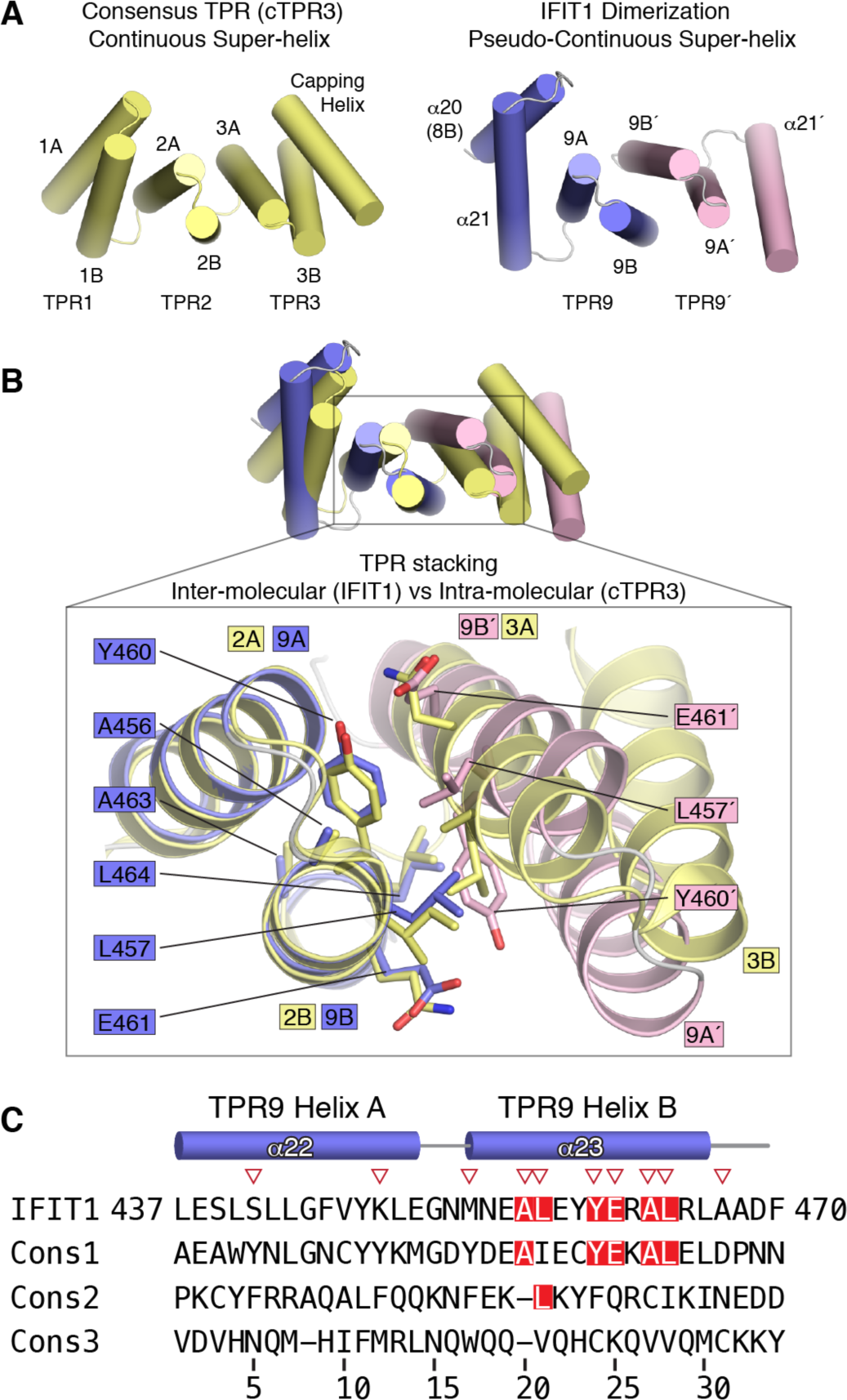
IFIT1 forms a pseudo-continuous TPR super-helix that mediates dimerization. **(A)** Left, structure of cTPR3 (PDB ID: 1NA0) made up of 3 consensus repeats, and Right, the central region from the IFIT1 dimer colored according to the molecule each helix originates from. Together, the C-terminal helices from each IFIT1 molecule associate in a manner that resembles a continuous TPR array. **(B)** Superposition of cTPR3 over the IFIT1 dimerization helices and close-up of residues involved in IFIT1 inter-molecular TPR stacking vs cTPR3 intra-molecular TPR stacking. **(C)** Comparison of IFIT1 TPR9 to consensus TPR sequences, which represent the top 3 residue preferences for each position of the 34 amino acid TPR motif. The consensus sequences were adapted from ref. (4).

### IFIT1 dimerization is dispensable for full RNA binding and inhibition of translation

To investigate the role of dimerization in RNA binding by IFIT1, we performed an EMSA utilizing a 43-nucleotide m7Gppp-RNA with a five-residue overhang at the 5′ end, which is optimal for IFIT1 binding (6). The capped RNA was fluorescently labelled at the 3′ end with pCp-Cy5, and binding to wild-type IFIT1 and the monomeric mutant IFIT1^DM-E^ was tested (**Fig. 7A**). Both proteins bound the capped RNA with the same apparent affinity of ~ 50-100 nM, suggesting that dimerization is not required for optimal RNA binding. Moreover, SYBR-Gold stained EMSAs with other unlabeled m7Gppp-RNAs, including two virus derived sequences, showed similar results (**Fig. S6**).

**Figure 7.**
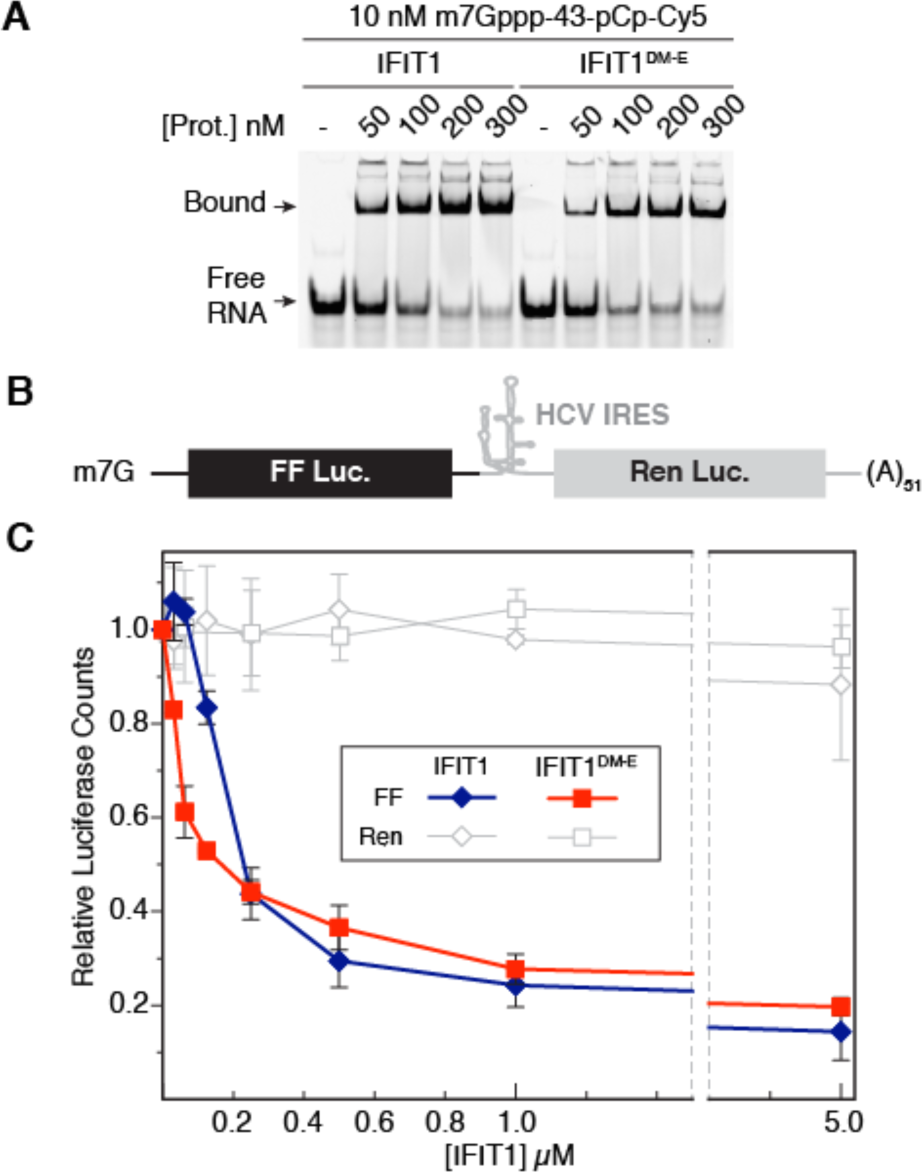
Functional analysis of IFIT1 dimerization. **(A)** Gel shift between wt IFIT1 or monomeric IFIT1^DM-E^ and a 43-nucleotide, capped-RNA which is fluorescently labelled with pCp-Cy5 at the 3′ end. **(B)** Schematic of bicistronic mRNA reporter utilized in the translation assay. **(C)** Comparison of translational inhibition by IFIT1 or IFIT1^DM-E^. Both proteins reduce cap-dependent FF translation equally without affecting cap-independent Ren translation. Data are represented as the mean of 2 independent measurements performed in duplicate ± standard deviation.

We next looked at the impact of dimerization on IFIT1 translational inhibition. To address this, we utilized an *in vitro* translation system consisting of Krebs extracts programmed with a bicistronic mRNA reporter (**Fig. 7B**) (30). The first open reading frame (ORF) produces Firefly luciferase (FF) in a cap-dependent manner, and should be susceptible to IFIT1 inhibition, while the second ORF expresses *Renilla* luciferase (Ren) under the control of an internal ribosome entry site (IRES) from hepatitis C virus (HCV), and would reveal the levels non-specific IFIT1 activity. Titrating IFIT1 or IFIT1^DM-E^ into the extracts at concentrations of 30 nM to 5 μM resulted in similar reductions of FF levels (compared to buffer control) (**Fig. 7C**). In both cases, Ren levels only dropped by 5-10 % indicating minimal non-specific activity. Thus, IFIT1 and IFIT1^DM-E^ exert similar translation inhibition, confirming that homo-dimerization is not required for IFIT1 inhibitory activity in an *in vitro* setting.

### RNA binding is associated with large-scale conformational changes

The means by which RNA gains entry to its binding site in IFIT1 is not immediately apparent, particularly because of the narrow shape and depth of the tunnel. To gain insight into the RNA-dependent conformational changes, we performed limited protease digestion of IFIT1 with trypsin and endoproteinase Glu-C, which revealed that both PPP-oligoA- and capped-oligoA-bound IFIT1 exhibit a greater degree of stabilization than RNA-free IFIT1 (**Fig. 8A**). The capped-RNA bound complex was more resistant to proteolysis than PPP-RNA-bound IFIT1, which reflects either specific conformational changes associated with capped-RNA binding or, more likely, the stronger affinity of IFIT1 towards capped-RNA (6, 22).

**Figure 8.**
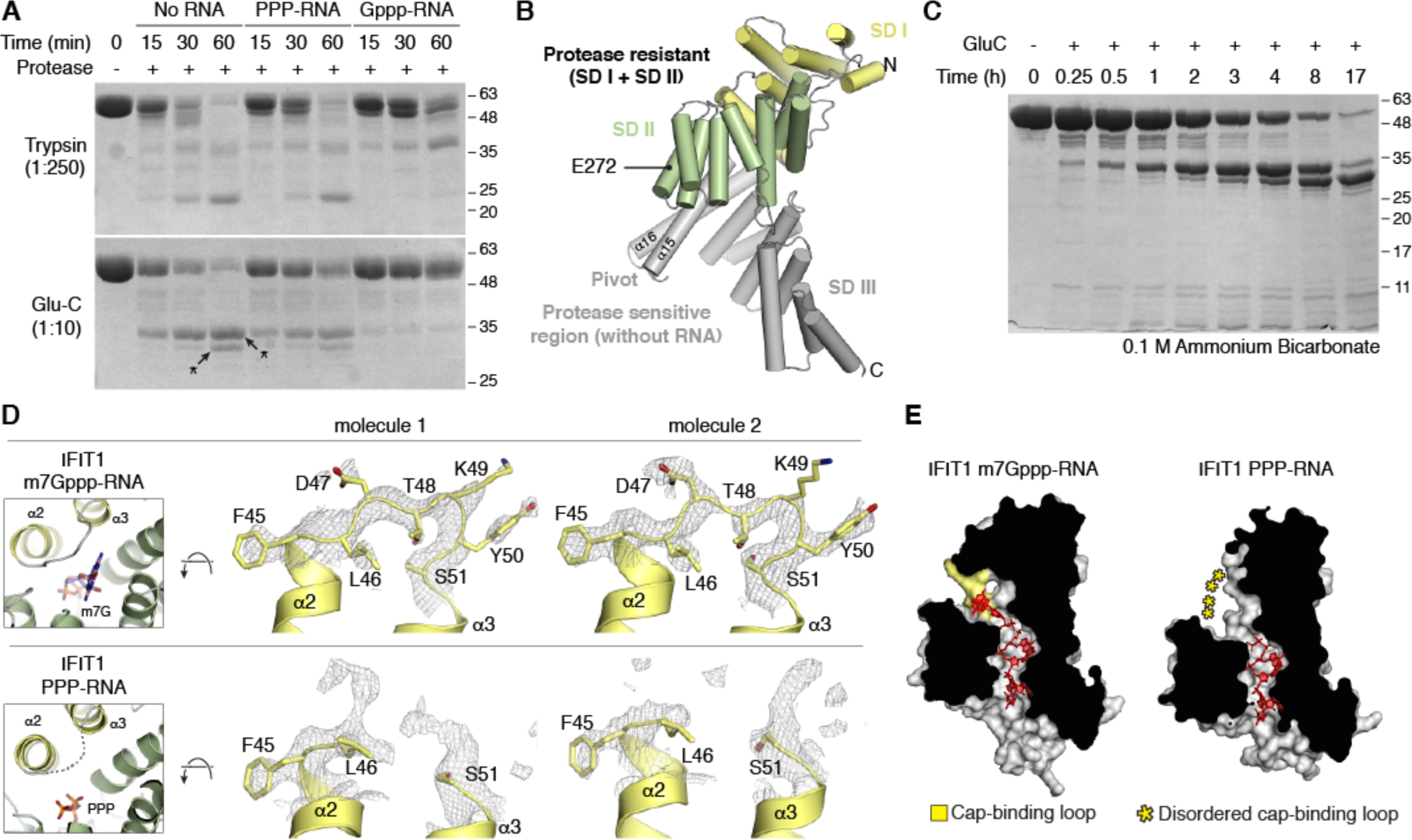
Limited protease digestion of IFIT1 and conformational changes associated with RNA binding. **(A)** IFIT1, IFIT1 with PPP-oligoA, and IFIT1 with Gppp-oligoA were subjected to limited protease digestion for the indicated times. Top, 1:250 (w/w) IFIT1:trypisin and Bottom, 1:10 (w/w) IFIT1:Endoproteinase Glu-C. In this assay, Gppp-RNA was used instead of m7Gppp-RNA since it has been shown to bind similarly (6). The two indicated bands were excised and analyzed by in-gel tryptic digestion and LC/MS/MS to map the in-solution Glu-C cleavage sites. **(B)** Location of Glu 272 on subdomain II, where Glu-C cleaves IFIT1 to produce the 32 kDa bottom fragment. **(C)** Extended time course of Glu-C proteolysis in 0.1M Ammonium Bicarbonate pH 7.8 with 1:10 protease:protein. **(D)** Simulated annealing 2F_o_-F_c_ omit map of IFIT1 residues 45- 51. The cap-binding loop is more ordered in the m7Gppp-RNA bound structure compared to the PPP-RNA bound structure. **(E)** Cross-section of the IFIT1 RNA-binding tunnel (grey protein surface with RNA depicted as red sticks). The cap-binding pocket is an open state in the absence of an m7G moiety.

The two major stable fragments resulting from endoproteinase Glu-C treatment of IFIT1 without RNA have apparent MWs of ~ 35 and ~ 32 kDa (**Fig. 8A, bottom**). We further examined these using in-gel tryptic digestion followed by LC/MS/MS, which showed that both originate from the protein N-terminus, namely subdomains I and II (**Fig. S7, A and B**). Intact protein MS of the second fragment indicates that it is the result of cleavage at Glu 272 (**Fig. S7, C and D**), and corresponds to the elastase-resistant fragment of human IFIT1 previously crystallized (**Fig. 8B**). When Glu-C cleavage was repeated in 0.1 M ammonium bicarbonate buffer, we observed slower kinetics of degradation, which we took advantage of to follow the IFIT1/Glu-C degradation profile over a period of 17 hours (**Fig. 8C**). The extended time course shows that 32 kDa fragment is derived from the ~ 35 kDa fragment, and also reveals the presence of several intermediate ~ 40- 48 kDa fragments within the first hour, which are likely due to cleavage at one of several positions on subdomain III. Thus, it would appear that the IFIT1 C-terminus exists in one or more open conformations that are susceptible to limited protease digestion, facilitating RNA entry into the narrow tunnel.

### A flexible cap-binding loop facilitates m7G recognition

Given that mRNA likely enters the tunnel from the C-terminal side of the protein, it is unclear how the m7G moiety is accommodated inside the cap-binding pocket, which forms a narrow, circular extension at the N-terminal side of the tunnel (**Fig. 2B**). To address this, we compared the structures of PPP-RNA-bound and m7Gppp-RNA-bound IFIT1, which revealed small-scale conformational within the cap-binding loop (residues 45-50). This loop forms one wall of the cap-binding pocket and is stabilized by binding of the m7G moiety, as it is somewhat disordered in the PPP-RNA structure (**Fig. 8, D and E**). However, based on crystal temperature factors it seems that the loop retains its dynamic nature even in the presence of the cap (**Fig. S8A**). In the crystal structure of RNA-free, N-terminal IFIT1 (composed of SD I+II (5)), the cap-binding loop also appears to be mobile as it has high temperature factors (**Fig. S8B**). Interestingly, these small-scale conformational changes were not evident from the high-resolution structures of the monomeric IFIT1^DM-E^ bound to PPP- and m7Gppp-RNA, since the cap-binding pocket of its PPP-RNA bound form was occupied by PEG molecules from the crystallization solution, artificially stabilizing the cap-binding loop (6). Nevertheless, the plasticity of the cap-binding loop is likely to be an important element in m7G recognition, as it would facilitate mRNA binding by leaving the cap-binding pocket in a more open and accessible state (**Fig. 8E**). On the other hand, the critical cap-binding residue Trp 147 is coordinated by Glu 176 in all structures, and appears to form a pre-organized ‘landing pad’ for m7G binding (**Fig. 2D**). Supporting the importance of the W147-E176 interaction for IFIT1 function, disrupting this coordinating pair through mutation (W147F or E176A) reduced binding to capped-RNA and translational inhibitory activity (6).

## Discussion

The Interferon-Induced Proteins with Tetratricopeptide Repeats family has been reported to mediate their antiviral effects through various mechanisms (31). Central to this ability is the TPR motif, a versatile protein-protein interaction module which in IFITs has also adapted to interact with virus-derived RNA (5, 6, 32). One facet of IFIT antiviral activity is the formation of an IFN-dependent, multi-protein complex with IFIT1, IFIT2, and IFIT3 at its core (19). The mechanisms regulating homo- and hetero-oligomerization of the IFIT core of this complex remain poorly characterized. The crystal structure of IFIT2 showed that it homo-dimerizes *via* domain swapping in which the N-terminal portion of its subdomain II (helices α7-α9) is exchanged between the interacting IFIT2 molecules (8). We report here the structural basis for IFIT1 dimerization, which utilizes a different mechanism for self-association. IFIT1 dimerizes through the formation of a C-terminal, pseudo-continuous TPR super-helix, which forms in a manner that mimics naturally occurring, intra-molecular TPR-stacking interactions that normally give rise to continuous TPR arrays.

TPR containing proteins have been shown to assemble into homo-dimers and higher order complexes using diverse mechanisms, ranging from those which resemble TPR-peptide interactions (e.g. the self-assembling, magnetosome-forming MamA (33)), to interactions which involve entire TPR domains encircling each other and burying up to 6000 Å^2^ (e.g. subunits of the anaphase-promoting complex (34)). The crystal structure of an engineered TPR domain made up of consensus motifs (CTPR3Y3) has been shown to assemble into trimers with one interface forming a C-terminal pseudo-continuous super-helix (35), similar to IFIT1. As with IFIT1, the pseudo-continuous super-helix of CTPR3Y3 involves displacement of a C-terminal capping helix, except in IFIT1 this capping helix appears to be lost due to a C-terminal truncation (when compared to IFIT5). Nevertheless, our structure confirms the notion that displacement or loss of a capping-helix giving rise to pseudo-continuous TPR arrays may be a general mechanism for TPR protein oligomerization (35). We propose that TPR9 can be used as a TPR dimerization module to facilitate the design of self-assembling TPR molecules. We demonstrated the feasibility of this by grafting the IFIT1 TPR9 sequence onto human IFIT5, thereby converting it from a monomer to a dimer.

TPR proteins have also been reported to form crystallographic dimers that wouldn’t otherwise form in solution (36), but we thoroughly validated the presence of IFIT1 dimerization in solution using gel-filtration, SEC-MALS, and AUC. These data are consistent with the formation of IFIT1 oligomers during RNA-binding gel shift assays and during protein blue-native PAGE electrophoresis (22). Additionally, human and rabbit IFIT1 were shown to bind capped-RNA with low-level positive cooperativity (Hill coefficient, n = 1.6-1.7) (22). However, co-immunoprecipitation assays failed to detect an IFIT1 self-interaction in human cells (19), possibly because the interaction is relatively weak. This is supported by our AUC analysis where we observed that wild-type IFIT1 had established a rapidly reversible monomer-dimer equilibrium during centrifugation. Furthermore, the relatively small dimerization interface burying only ~ 530 Å^2^ is smaller than expected for a stable self-interaction (37).

Regardless, during AUC IFIT1 is still predominantly dimeric at 3.3 μM, and given that IFIT1 is highly expressed upon induction where its levels can approach 1-2 million copies per cell (19), it is likely that physiological concentrations of IFIT1 are within a range that promotes dimer formation. Moreover, IFIT1 has been shown to re-localize to discrete spots following transfection of PPP-RNA (19), where even higher local concentrations would favor dimerization. However, although IFIT1 dimer formation *in vivo* is a likely scenario, the possibility remains that it is non-physiological. Since human IFIT1 is at the center of a large multi-protein complex (19), isolating it *in vitro* could give rise to artifacts. In this case, IFIT1 homo-dimerization may be due to residual binding that is a consequence of IFIT1 hetero-oligomerization with other members of the interactome, such as IFIT2 or IFIT3.

Most mammals encode an IFIT1 protein and an IFIT1B protein (3). However, in mice, IFIT1 appears to be deleted and instead they have three copies of IFIT1B (3), referred to as mouse Ifit1, mouse Ifit1b, and mouse Ifit1c (with 58-77 % sequence identity between them). On the other hand, in humans, IFIT1B expression has not been shown to change with IFN treatment (27), and heterologous overexpression in yeast suggests that it lacks a function in RNA binding and translational inhibition (3). Nevertheless, the conserved nature of the IFIT1 dimerization interface suggests that mammalian IFIT1B proteins homo-dimerize in the same manner, and may even form hetero-dimers with IFIT1 or other IFIT1B-like proteins through the same interface. Supporting this notion, mouse Ifit1c was found to co-purify with mouse Ifit1 in capped-RNA pulldowns (21). Although it remains to be shown, hetero-dimerization through this interface could regulate RNA binding or serve to expand the protein-protein interaction network of IFIT1, therefore diversifying IFIT1-mediated antiviral responses.

Although foot-printing primer-extension assays performed under equilibrium binding conditions revealed that human IFIT1 can bind capped-RNA with positive cooperativity (22), our gel-shift RNA binding assays did not reveal a difference in apparent affinity between monomeric and dimeric IFIT1. One possible explanation for this discrepancy is that EMSAs are not a true equilibrium method, and complex formation is susceptible to variations in the assay conditions (38, 39), therefore the apparent RNA-binding affinities of IFIT1 and the monomeric mutant could be affected. Future assays performed under equilibrium conditions (e.g. primer-extension foot-printing, fluorescence polarization, or isothermal titration calorimetry) could be performed to clarify the role of dimerization in RNA binding. Of note, our attempts at isothermal titration calorimetry with human IFIT1 were unsuccessful due protein precipitation during the experiment. Regardless, the translation assays in Krebs extracts indicate that homo-dimerization is dispensable for translational inhibition by IFIT1, at least under this *in vitro* setting.

Finally, comparison of the different RNA bound forms of IFIT1 complemented with limited protease digestion revealed large-scale and small-scale conformational changes associated with RNA binding. Based on the structural analysis, limited proteolysis, and previous work with human IFIT5 (5), we propose the following model for capped mRNA binding by IFIT1 in solution. IFIT1 exists in one or more open conformations that are amenable for RNA entry into the tunnel. Upon binding the 5′ end of mRNA (nucleotides 1-4), IFIT1 undergoes conformational changes at its C-terminus that are facilitated by the pivot region, as was seen in IFIT5 (5). At the same time, or in a subsequent step, the m7G moiety is directed towards the cap-binding pocket, which would initially be in a more open and accessible state as indicated by the dynamic nature of the cap-binding loop. Recognition by Trp 147 and stabilization of the cap-binding loop would then be the last in a series of coordinated steps required for mRNA binding. The RNA-dependent conformational changes would help IFIT1 clamp down on viral mRNA, thus decreasing its rate of dissociation, and prevent translation initiation factors from outcompeting IFIT1 once it is bound. Both the large scale and small scale conformational changes described here are also likely impacted by other members of the IFIT interactome, further enhancing or limiting IFIT1 antiviral activity *in vivo*.

In summary, we have determined the crystal structure of human IFIT1, a central component of the IFIT interactome, in its native, dimeric form. The structure shows that IFIT1 uses a small C-terminal interface to form extended, antiparallel dimers that are distinct from the parallel, domain-swapped dimers of IFIT2. This arrangement of IFIT1 dimers could combine with the parallel interface found in IFIT2 to initiate interactome assembly. Alternatively, IFIT2 or IFIT3 could compete with the IFIT1 homo-dimerization interface to build up the interactome. As these interactions could be important for regulating IFIT1 conformational changes and inhibitory activity, our results will contribute to piecing together the network of interactions that regulate IFIT proteins’ role in the antiviral innate immune response.

### Experimental Procedures

#### Plasmids

IFIT1, IFIT2, and IFIT5 were sub-cloned into pSMT3 between BamHI and NotI sites, resulting in a fusion protein with an N-terminal, Ulp1-cleavable 6xHis-Sumo tag (40). IFIT1 point mutants were generated by site-directed mutagenesis using iProof High-Fidelity DNA polymerase (Bio-Rad). The chimera made up of IFIT5 (residues 1-435) and IFIT1 (residues 437-478) was generated using overlap-extension PCR, and sub-cloned into pSMT3 between BamHI and NotI.

#### Expression and purification of IFIT proteins

IFIT1, IFIT1 mutants and IFIT5 expression and purification are described elsewhere (6). The IFIT5/IFIT1 chimera was expressed in Rosetta 2 (DE3) pLysS BL21 cells (Novagen) by first growing transformed cells in LB at 37 °C until an OD600 of 0.6-0.8, followed by induction with 1 mM IPTG and growing the cells for 4 hours at 30 °C. The protein was purified by Ni-affinity chromatography followed by cleavage of the tag. The cleaved protein was further purified by passing over a 5 ml HiTrap SP HP column (GE Healthcare) equilibrated with 50 mM HEPES pH 7.5 and 1 mM DTT, followed by washing with 50 mM HEPES pH 7.5, 100 mM NaCl, and 1 mM DTT, and eluted with 50 mM HEPES pH 7.5, 400 mM NaCl, and 1 mM DTT. IFIT2 was expressed in BL21 (DE3) and purified using 2-step Ni-affinity chromatography (with cleavage of the tag) using standard protocols. All proteins were further purified by gel filtration using superdex 200 10/300 or 16/60 columns (GE Healthcare) in 20 mM Tris pH 7.6, 150 mM NaCl and 1 mM DTT.

#### Preparation of PPP and m7Gppp containing oligoadenylate RNA

PPP-AAAA and Gppp-AAAA were prepared using phosphoramidite solid phase synthesis followed by installation of the triphosphate or cap moieties as previously described (6); the oligos were purified by HPLC and characterized by LC-MS as before (6). m7Gppp-AAAA was prepared by enzymatically modifying Gppp-AAAA with purified mRNA cap guanine-N7 methyltransferase as before (6).

#### Crystallization and data collection

Wild-type human IFIT1 was mixed with molar excess chemically synthesized oligos either directly before crystallization or prior to gel filtration. The protein buffer was 20 mM Tris pH 7.6, 150 mM NaCl and 1 mM TCEP. The co-crystals were obtained with 2-4 mg/ml protein drops mixed 1:1 with reservoir solution containing 27-32 % PEG 200 (Sigma), 0.1 M Tris pH 8.1, and 200 mM CaCl2 using the hanging drop vapour diffusion method at 4 °C. Crystals were flash frozen in liquid nitrogen without additional cryo-protection. All data were collected at 100K with 0.979 Å X-rays on beamline 08ID-1 at the CLS, which was equipped with a Mar300CCD detector (41), and the images integrated, scaled, and merged using the HKL2000 suite (43). Data were truncated with Ctruncate (ccp4 suite (42)) and 5 % of reflections were set aside for the R_free_ set. Only the highest resolution dataset was used as a master file to generate all R_free_ sets. Data collection statistics from HKL2000 are in **Table S2**.

#### Structure determination, model building, and refinement

The initial crystal structure was determined by molecular replacement using the PHASER program in Phenix (43). The structure of RNA-bound human IFIT5 (residues 1-434) was used as a search model. The remaining 3 helices of IFIT1 (residues 437-469) were built manually after one macro-cycle of refinement in Phenix (45). Subsequent structures were determined with rigid body refinement with the protein only. Restraints for m7GpppA were calculated using Phenix eLBOW (44). The structures were refined iteratively using Phenix with manual model building in Coot (46). The refinement strategy included all-isotropic B-factor refinement and TLS. The final models contained protein residues 9-84, 91-305, and 310-469 for m7Gppp-RNA bound IFIT1 chain A; 9-84, 90-304, and 315-469 for m7Gppp-RNA bound IFIT1 chain C; 9-46, 51-81, 92-247, 251-301, and 316-469 for PPP-RNA bound IFIT1 chain A; and 9-46, 51-81, 92-247, 251-305, and 314-469 for PPP-RNA bound IFIT1 chain C. Structure validation was performed with MOLPROBITY in Phenix (47). Ramachandran statistics are as follows: IFIT1 with m7Gppp-RNA, 98.5 % favored, 0 % outliers, 0.25 % rotamer outliers; and IFIT1 with PPP-RNA: 98.2 % favored, 0 % outliers, 0.4 % rotamer outliers. The MOLPROBITY overall scores for the m7Gppp-RNA and PPP-RNA-bound structures were 0.58 and 0.63, respectively. Refinement statistics are in **Table S2**.

#### Sequence and structure analysis

APBS was used to calculate the surface electrostatic potential (48), and PyMol to general all molecular figures (https://www.pymol.org/). For surface electrostatic analysis, all surfaces are colored by electrostatic potential from negative (-10 kTe-1; red) to positive (+10 kTe-1; blue). ESPript was used to generate the sequence alignment in Fig. 4A (49). Sequence conservation analysis by WebLogo (50) in Fig. 4B was performed as described (6).

#### Analytical gel filtration

500 μL (Fig. 5A) or 250 μL (Fig. 5B) of purified protein at 2 mg/ml were injected onto a superdex 200 10/300 increase column (GE Healthcare) equilibrated in 20 mM Tris pH 7.6 and 150 mM NaCl. 1 μg (Fig. 5A) or 2 μg (Fig. 5B) of the input proteins were run on 12 % SDS-PAGE and visualized by coomassie staining.

#### Analytical ultracentrifugation

Sedimentation velocity experiments were performed in a Beckman XL-I analytical ultracentrifuge using an An-60Ti rotor. Solutions containing 20 mM Tris pH 7.6, 150 mM NaCl, 0.5 mM TCEP and varying concentrations of IFIT1 (3.3 μM, 8.5 μM , and 18.0 μM) were centrifuged at 50,000 rpm over approximately 8h. A total of 200 scans were collected per run, with absorbances measured at 280 nm. Sedimentation velocity data were fit to the *c*(*S*) model using the computer program SEDFIT (28). Sedimentation coefficients were determined by integration of c(S) distributions in SEDFIT and then transformed into *S*_20,W_ values. The program SEDNTERP (v1.09, http://www.jphilo.mailway.com/) was used for calculation of solution viscosity, solution density, and IFIT1 partial specific volume (from primary amino acid sequence) at each temperature examined.

#### Multi-angle light scattering with inline size-exclusion chromatography (SEC-MALS)

50 μL of purified IFIT1 (10 mg/ml), IFIT1 ^DM-E^ (12 mg/ml), or BSA (10.5 mg/ml) were injected onto superdex 200 10/300 increase (GE Healthcare) equilibrated with 20 mM Tris pH 7.6 and 150 mM NaCl. Measurements were performed using a Waters e2796 Separations module (Waters Corporation) equipped with an in-line Waters 2489 UV/Visible detector set to 280nm (Waters corporation), a miniDAWN TREOS multi-angle light scattering detector with a 656nm laser (Wyatt Technology), and an Optilab rEX differential refractive index detector (Wyatt Technology). Data was processed and analyzed using the ASTRA software package (version 5.3.4.18 from Wyatt technology).

#### Limited Protease Digestion

PPP-oligoA or Gppp-oligoA were re-suspended in water at a final concentration of 10 nmol/μL. 1.26 nmol IFIT1 (70 μg) in gel filtration buffer were mixed with either 15 μL water, 15 μL PPP-oligoA (1.5 nmol), or 15 μL Gppp-oligoA (1.5 nmol) at a final volume of 70 μL, resulting in 10 mg/ml (10X) protein stocks with or without RNA. 10X protease stocks were prepared in 20 mM HEPES pH 7.5, 50 mM KCl, and 20 mM Mg_2_SO_4_ (Cleavage buffer), resulting in a 1 mg/ml endoproteinase Glu-C stock and 0.04 mg/ml trypsin stock. For each reaction (corresponding to each lane in Fig. 8A), 1.6 μL of 10X protein and 1.6 μL of 10X protease were mixed at a final volume of 16 μL in Cleavage buffer and the reaction carried out at room temperature. At the indicated time points, the reactions were stopped with 4 μL of 5X SDS loading buffer, boiled for 10 minutes, and stored at -20 °C until the samples were ready to be run on 12 % SDS-PAGE. For the extended time-course (also performed at room temperature), 150 μg IFIT1 was mixed with 15 μg Glu-C in 150 μL containing 0.1 ammonium bicarbonate buffer pH 7.8. At the indicated time points, 16 μL of the reaction were removed and mixed with 4 μL of 5X SDS loading buffer, then boiled for 10 minutes, and stored at -20 °C until the samples were ready to be run on 12 % SDS-PAGE.

#### RNA electro-mobility gel shift assays (EMSA)

Capped RNAs for EMSAs were prepared by *in vitro* transcription with T7 RNA polymerase followed by PAGE purification and post-transcriptional capping with the vaccinia capping system (NEB) as previously described (6). Preparation of pCp-labelled RNA was performed with T4 RNA ligase (NEB) as previously described (6). Gel shifts were performed exactly as described before (6).

#### Circular dichroism (CD) spectroscopy

IFIT1, IFIT1^DM-A^, or IFIT1^DM-E^ were buffer exchanged into 50 mM Na_2_HPO_4_/NaH_2_PO_4_ pH 8 and diluted to a final concentration of 0.5 mg/ml. Measurements were performed on a Chirascan CD spectrometer using a 0.2 mm quartz cell. CD wavelength scans were obtained at 21 °C, with 1 nm step size and 0.5 s time intervals. The signals obtained were in CD millidegrees and are represented as an average of three spectra obtained for each sample. The averaged spectra were smoothened using the Savitsky-Golay method and a window size of 3.

#### In vitro translation assay with Krebs extracts

Preparation of reporter mRNA and *in vitro* translation assay were performed as before (6). Briefly, pSP-(CAGless)/FF/HCV/Ren plasmid was linearized with BamHI and transcribed with SP6 RNA polymerase in the presence of m7G(5′)ppp(5′)G cap analog (NEB). *In vitro* translations were set up at a final volume of 10 μl with 4 ng/*μ*L reporter mRNA (~ 4 nM final) and 1 μL wild-type IFIT1 or IFIT1^DM-E^ in untreated Krebs-2 extracts. The reactions were set up on ice and translation was allowed to proceed for 1 hr at 30 °C. The reactions were stopped on ice, and FF and Ren luciferase activities (RLU) were measured on a Berthold Lumat LB 9507 luminometer. Values obtained were normalized against buffer control, which was set at 1. The following final protein concentrations were used: 0.031, 0.062, 0.125, 0.250, 0.5, 1, and 5 μM.

## Accession codes

Coordinates and structure factor data for the structures reported in this study were deposited in the Protein Data Bank under the accession codes 5W5H (IFIT1 + m7Gppp-AAAA) and 5W5I (IFIT1 + PPP-AAAA)

## Acknowledgements

We thank the CMCF staff for X-ray data collection performed on beamline 08ID-1 at the Canadian Light Source, which is supported by the Canada Foundation for Innovation, Natural Sciences and Engineering Research Council of Canada, the University of Saskatchewan, the Government of Saskatchewan, Western Economic Diversification Canada, the National Research Council Canada, and the Canadian Institutes of Health Research. Analytical ultracentrifugation experiments were performed using a Beckman XL-I analytical ultracentrifuge housed in the Concordia Centre for Structural and Functional Genomics (CSFG). We thank Piratip Pratumsuwan and Stefannie Neun for preliminary protein characterization and technical assistance, and Maxime Isabelle for protein mass spectrometry. Support was provided by the CIHR Strategic Training Initiative in Chemical Biology and the NSERC CREATE Training Program in Bionanomachines (Y.M.A.); a Canada Research Chair (B.N.); Canadian Institutes of Health Research Grants (MOP-133535, B.N. and FDN-148366, J.P.); Discovery Grants from the Natural Sciences and Engineering Research Council of Canada (M.J.D. and P.D.P.); and the McGill CIHR Drug Development Training Program (S.M-M.).

## Conflict of interest

The authors declare that they have no conflicts of interest with the contents of this article.

## Author Contributions

B.N. and Y.M.A. designed the study and prepared the manuscript with input from all authors; B.N., J.P., and M.J.D. supervised the work; Y.M.A. prepared proteins and RNA, and performed structural work, SEC-MALS, limited proteolysis, CD spectroscopy, analytical gel filtration, and EMSAs; S.M-M. performed chemical synthesis and purification of PPP- and Gppp-oligoA; R.C. prepared translation extracts and reporter mRNA, and performed translation assays; P.D.P. performed AUC data collection and analysis.

## References

1. Fensterl, V., Chattopadhyay, S., and Sen, G. C. (2015) No Love Lost Between Viruses and Interferons. Annual Review of Virology. 2, 549–572

2. Fensterl, V., and Sen, G. C. (2015) Interferon-induced Ifit proteins: their role in viral pathogenesis. J Virol. 89, 2462–2468

3. Daugherty, M. D., Schaller, A. M., Geballe, A. P., and Malik, H. S. (2016) Evolution-guided functional analyses reveal diverse antiviral specificities encoded by IFIT1 genes in mammals. elife. 5, 60

4. Main, E. R. G., Xiong, Y., Cocco, M. J., D'Andrea, L., and Regan, L. (2003) Design of stable alpha-helical arrays from an idealized TPR motif. Structure. 11, 497–508

5. Abbas, Y. M., Pichlmair, A., Górna, M. W., Superti-Furga, G., and Nagar, B. (2013) Structural basis for viral 5′-PPP-RNA recognition by human IFIT proteins. Nature. 494, 60–64

6. Abbas, Y. M., Laudenbach, B. T., Martínez-Montero, S., Cencic, R., Habjan, M., Pichlmair, A., Damha, M. J., Pelletier, J., and Nagar, B. (2017) Structure of human IFIT1 with capped RNA reveals adaptable mRNA binding and mechanisms for sensing N1 and N2 ribose 2′-O methylations. Proceedings of the National Academy of Sciences. 10.1073/pnas.1612444114

7. Katibah, G. E., Lee, H. J., Huizar, J. P., Vogan, J. M., Alber, T., and Collins, K. (2013) tRNA binding, structure, and localization of the human interferon-induced protein IFIT5. Mol Cell. 49, 743–750

8. Yang, Z., Liang, H., Zhou, Q., Li, Y., Chen, H., Ye, W., Chen, D., Fleming, J., Shu, H., and Liu, Y. (2012) Crystal structure of ISG54 reveals a novel RNA binding structure and potential functional mechanisms. Cell Res. 22, 1328–1338

9. Feng, F., Yuan, L., Wang, Y. E., Crowley, C., Lv, Z., Li, J., Liu, Y., Cheng, G., Zeng, S., and Liang, H. (2013) Crystal structure and nucleotide selectivity of human IFIT5/ISG58. Cell Res. 23, 1055–1058

10. D’Andrea, L. D., and Regan, L. (2003) TPR proteins: the versatile helix. Trends Biochem Sci. 28, 655–662

11. Feng, X., Wang, Y., Ma, Z., Yang, R., Liang, S., Zhang, M., Song, S., Li, S., Liu, G., Fan, D., and Gao, S. (2014) MicroRNA-645, up-regulated in human adencarcinoma of gastric esophageal junction, inhibits apoptosis by targeting tumor suppressor IFIT2. BMC Cancer. 14, 633

12. Xiao, S., Li, D., Zhu, H.-Q., Song, M.-G., Pan, X.-R., Jia, P.-M., Peng, L.-L., Dou, A.-X., Chen, G.-Q., Chen, S.-J., Chen, Z., and Tong, J.-H. (2006) RIG-G as a key mediator of the antiproliferative activity of interferon-related pathways through enhancing p21 and p27 proteins. Proc Natl Acad Sci USA. 103, 16448–16453

13. Lai, K. C., Liu, C. J., Chang, K. W., and Lee, T. C. (2013) Depleting IFIT2 mediates atypical PKC signaling to enhance the migration and metastatic activity of oral squamous cell carcinoma cells. Oncogene. 32, 3686–3697

14. Lai, K.-C., Chang, K.-W., Liu, C.-J., Kao, S.-Y., and Lee, T.-C. (2008) IFN-induced protein with tetratricopeptide repeats 2 inhibits migration activity and increases survival of oral squamous cell carcinoma. Mol. Cancer Res. 6, 1431–1439

15. Stawowczyk, M., Van Scoy, S., Kumar, K. P., and Reich, N. C. (2011) The interferon stimulated gene 54 promotes apoptosis. Journal of Biological Chemistry. 286, 7257–7266

16. Berchtold, S., Manncke, B., Klenk, J., Geisel, J., Autenrieth, I. B., and Bohn, E. (2008) Forced IFIT-2 expression represses LPS induced TNF-alpha expression at posttranscriptional levels. BMC Immunol. 9, 75

17. Li, Y., Li, C., Xue, P., Zhong, B., Mao, A.-P., Ran, Y., Chen, H., Wang, Y.-Y., Yang, F., and Shu, H.-B. (2009) ISG56 is a negative-feedback regulator of virus-triggered signaling and cellular antiviral response. Proceedings of the National Academy of Sciences. 106, 7945–7950

18. Liu, X. Y., Chen, W., Wei, B., Shan, Y. F., and Wang, C. (2011) IFN-Induced TPR Protein IFIT3 Potentiates Antiviral Signaling by Bridging MAVS and TBK1. The Journal of Immunology. 187, 2559–2568

19. Pichlmair, A., Lassnig, C., Eberle, C.-A., Górna, M. W., Baumann, C. L., Burkard, T. R., Bürckstümmer, T., Stefanovic, A., Krieger, S., Bennett, K. L., Rülicke, T., Weber, F., Colinge, J., Müller, M., and Superti-Furga, G. (2011) IFIT1 is an antiviral protein that recognizes 5′-triphosphate RNA. Nat Immunol. 12, 624–630

20. Kimura, T., Katoh, H., Kayama, H., Saiga, H., Okuyama, M., Okamoto, T., Umemoto, E., Matsuura, Y., Yamamoto, M., and Takeda, K. (2013) Ifit1 inhibits JEV replication through binding to 5′ capped 2′-O unmethylated RNA. J Virol. 87, 9997–10003

21. Habjan, M., Hubel, P., Lacerda, L., Benda, C., Holze, C., Eberl, C. H., Mann, A., Kindler, E., Gil-Cruz, C., Ziebuhr, J., Thiel, V., and Pichlmair, A. (2013) Sequestration by IFIT1 Impairs Translation of 2′O-unmethylated Capped RNA. PLoS Pathog. 9, e 1003663

22. Kumar, P., Sweeney, T. R., Skabkin, M. A., Skabkina, O. V., Hellen, C. U. T., and Pestova, T. V. (2014) Inhibition of translation by IFIT family members is determined by their ability to interact selectively with the 5“-terminal regions of cap0-, cap1- and 5”ppp-mRNAs. Nucleic Acids Res. 42, 3228–3245

23. Hyde, J. L., Gardner, C. L., Kimura, T., White, J. P., Liu, G., Trobaugh, D. W., Huang, C., Tonelli, M., Paessler, S., Takeda, K., Klimstra, W. B., Amarasinghe, G. K., and Diamond, M. S. (2014) A viral RNA structural element alters host recognition of nonself RNA. Science. 343, 783–787

24. Daffis, S., Szretter, K. J., Schriewer, J., Li, J., Youn, S., Errett, J., Lin, T.-Y., Schneller, S., Zust, R., Dong, H., Thiel, V., Sen, G. C., Fensterl, V., Klimstra, W. B., Pierson, T. C., Buller, R. M., Gale, M., Jr, Shi, P.-Y., and Diamond, M. S. (2010) 2′-O methylation of the viral mRNA cap evades host restriction by IFIT family members. Nature. 468, 452–456

25. Zust, R., Cervantes-Barragan, L., Habjan, M., Maier, R., Neuman, B. W., Ziebuhr, J., Szretter, K. J., Baker, S. C., Barchet, W., Diamond, M. S., Siddell, S. G., Ludewig, B., and Thiel, V. (2011) Ribose 2′-O-methylation provides a molecular signature for the distinction of self and non-self mRNA dependent on the RNA sensor Mda5. Nat Immunol. 12, 137–143

26. Diamond, M. S. (2014) IFIT1: A dual sensor and effector molecule that detects non-2′-O methylated viral RNA and inhibits its translation. Cytokine Growth Factor Rev. 10.1016/j.cytogfr.2014.05.002

27. Fensterl, V., and Sen, G. C. (2011) The ISG56/IFIT1 gene family. Journal of Interferon & Cytokine Research. 31, 71–78

28. Schuck, P., and Balbo, A. (2005) Analytical Ultracentrifugation in the Study of Protein Self-association and Heterogeneous Protein-Protein Interactions. Cold Spring Harbor Laboratory Press

29. Krissinel, E., and Henrick, K. (2007) Inference of Macromolecular Assemblies from Crystalline State. J Mol Biol. 372, 774–797

30. Novac, O., Guenier, A. S., and Pelletier, J. (2004) Inhibitors of protein synthesis identified by a high throughput multiplexed translation screen. Nucleic Acids Res. 32, 902–915

31. Diamond, M. S., and Farzan, M. (2013) The broad-spectrum antiviral functions of IFIT and IFITM proteins. Nat Rev Immunol. 13, 46–57

32. Pichlmair, A., Schulz, O., Tan, C. P., Rehwinkel, J., Kato, H., Takeuchi, O., Akira, S., Way, M., Schiavo, G., and Reis e Sousa, C. (2009) Activation of MDA5 Requires Higher-Order RNA Structures Generated during Virus Infection. J Virol. 83, 10761–10769

33. Zeytuni, N., Ozyamak, E., Ben-Harush, K., Davidov, G., Levin, M., Gat, Y., Moyal, T., Brik, A., Komeili, A., and Zarivach, R. (2011) Self-recognition mechanism of MamA, a magnetosome-associated TPR-containing protein, promotes complex assembly. Proceedings of the National Academy of Sciences. 108, E480–7

34. Zhang, Z., Chang, L., Yang, J., Conin, N., Kulkarni, K., and Barford, D. (2013) The Four Canonical TPR Subunits of Human APC/C Form Related Homo-Dimeric Structures and Stack in Parallel to Form a TPR Suprahelix. J Mol Biol. 425, 4236–4248

35. Krachler, A. M., Sharma, A., and Kleanthous, C. (2010) Self-association of TPR domains: Lessons learned from a designed, consensus-based TPR oligomer. Proteins. 10.1002/prot.22726

36. Kajander, T., Cortajarena, A. L., Mochrie, S., and Regan, L. (2007) Structure and stability of designed TPR protein superhelices: unusual crystal packing and implications for natural TPR proteins. Acta Crystallogr D Biol Crystallogr. 63, 800–811

37. Conte, Lo, L., Chothia, C., and Janin, J. (1999) The atomic structure of protein-protein recognition sites. J Mol Biol. 285, 2177–2198

38. Gaudreault, M., Gingras, M.-E., Lessard, M., Leclerc, S., and Guérin, S. L. (2009) Electrophoretic Mobility Shift Assays for the Analysis of DNA-Protein Interactions. in Nucleic Acid Crystallography (Ennifar, E. ed), pp. 15–35, Methods in Molecular Biology™, Humana Press, Totowa, NJ, 543, 15–35

39. Fried, M. G., and Liu, G. (1994) Molecular sequestration stabilizes CAP–DNA complexes during polyacrylamide gel electrophoresis. Nucleic Acids Res. 22, 5054–5059

40. Mossessova, E., and Lima, C. D. (2000) Ulp1-SUMO crystal structure and genetic analysis reveal conserved interactions and a regulatory element essential for cell growth in yeast. Mol Cell. 5, 865–876

41. Grochulski, P., Fodje, M. N., Gorin, J., Labiuk, S. L., Berg, R., IUCr (2011) Beamline 08ID-1, the prime beamline of the Canadian Macromolecular Crystallography Facility. J Synchrotron Radiat. 18, 681–684

42. Winn, M. D., Ballard, C. C., Cowtan, K. D., Dodson, E. J., Emsley, P., Evans, P. R., Keegan, R. M., Krissinel, E. B., Leslie, A. G. W., McCoy, A., McNicholas, S. J., Murshudov, G. N., Pannu, N. S., Potterton, E. A., Powell, H. R., Read, R. J., Vagin, A., Wilson, K. S., IUCr (2011) Overview of the CCP4 suite and current developments. Acta Crystallogr D Biol Crystallogr. 67, 235–242

43. McCoy, A. J., Grosse-Kunstleve, R. W., Adams, P. D., Winn, M. D., Storoni, L. C., Read, R. J., IUCr (2007) Phaser crystallographic software. J Appl Crystallogr. 40, 658–674

44. Moriarty, N. W., Grosse-Kunstleve, R. W., and Adams, P. D. (2009) electronic Ligand Builder and Optimization Workbench (eLBOW): a tool for ligand coordinate and restraint generation. Acta Crystallogr D Biol Crystallogr. 65, 1074–1080

45. Adams, P. D., Afonine, P. V., Bunkóczi, G., Chen, V. B., Davis, I. W., Echols, N., Headd, J. J., Hung, L.-W., Kapral, G. J., Grosse-Kunstleve, R. W., McCoy, A. J., Moriarty, N. W., Oeffner, R., Read, R. J., Richardson, D. C., Richardson, J. S., Terwilliger, T. C., and Zwart, P. H. (2010) PHENIX: a comprehensive Python-based system for macromolecular structure solution. Acta Crystallogr D Biol Crystallogr. 66, 213–221

46. Emsley, P., Lohkamp, B., Scott, W. G., and Cowtan, K. (2010) Features and development of Coot. Acta Crystallogr D Biol Crystallogr. 66, 486–501

47. Chen, V. B., Arendall, W. B., Headd, J. J., Keedy, D. A., Immormino, R. M., Kapral, G. J., Murray, L. W., Richardson, J. S., Richardson, D. C., IUCr (2010) MolProbity: all-atom structure validation for macromolecular crystallography. Acta Crystallogr D Biol Crystallogr. 66, 12–21

48. Baker, N. A., Sept, D., Joseph, S., Holst, M. J., and McCammon, J. A. (2001) Electrostatics of nanosystems: Application to microtubules and the ribosome. Proceedings of the National Academy of Sciences. 98, 10037–10041

49. Robert, X., and Gouet, P. (2014) Deciphering key features in protein structures with the new ENDscript server. Nucleic Acids Res. 42, W320–4

50. Schneider, T. D., and Stephens, R. M. (2004) Sequence logos: a new way to display consensus sequences. Nucleic Acids Res. 18, 1–4

